# Multi-tissue methylation clocks for age estimation in the common bottlenose dolphin

**DOI:** 10.1101/2021.05.03.442523

**Authors:** Todd R. Robeck, Zhe Fei, Amin Haghani, Joseph A. Zoller, Caesar Z. Li, Karen J. Steinman, Stacy DiRocco, Lydia Staggs, Todd Schmitt, Steve Osborn, Gisele Montano, Magdalena Rodriguez, Steve Horvath

## Abstract

Accurate identification of individual ages within wild bottlenose dolphins (*Tursiops truncatus*) is critical for determining population health and the development of population management strategies. As such, we analyzed DNA methylation patterns by applying a custom methylation array (HorvathMammalMethyl40) to both blood (n = 140) and skin samples (n = 87) from known age or approximate age (0 to 57 years) bottlenose dolphins. We present three bottlenose dolphin specific age estimation clocks using combined blood and skin (48 CpGs, R = 0.93, median absolute error = 2.13 years), blood only (64 CpGs, R = 0.97, error= 1.46 years) and skin only (39 CpGs, R = 0.95, error= 2.53). Our sex estimator based on 71 CpGs predicts the sex of any odontocete species with 99.5% accuracy. We characterize individual cytosines that correlate with sex and age in dolphins.

The presented epigenetic clocks are expected to be useful for conservation efforts and for studying anthropogenic events.

## INTRODUCTION

Accurate age estimation of wild cetaceans is an important component of any population health assessment and is critical for the development of management plans designed to help wild populations in need[1]. For bottlenose dolphins, standard age estimations of wild animals rely on a combination of techniques including animal length (allometry), long term “capture and recapture” of animals using photo identification, and the counting of tooth growth layer groups (GLGs). Age estimators based on allometry lose accuracy once physical maturity has been reached and surpassed[2, 3]. Although long term photo identification studies have produced the most robust and accurate data concerning life history of a few populations of bottlenose dolphins, these studies face limitations, e.g. the inability to accurately define animals that had already reached physical maturity at the start of the surveys[4, 5]. Although long-term photo identification programs have been used with a few populations of bottlenose dolphins, their high economic costs and long-term time commitments render them less effective for rapid and time-sensitive evaluations of critically endangered or threatened populations.

Ages of odontocetes can be estimated based on tooth growth layer groups (GLGs[6]). Although GLGs represent the current gold standard for determining the age of bottlenose dolphins, the requirement for a tooth may render this method unacceptably invasive for many research efforts. Finally, some debate exists concerning the accuracy of relying on growth ring deposition for age estimation older animals that may have experienced significant tooth wear[7].

As such, other aging methods have been evaluated for use with bottlenose dolphins or other cetaceans including eye lens aspartic acid racemization[8, 9], fatty acid composition[10–12], radiocarbon 14 dating from fallout[13], telomere length[14], radiograph changes in tooth to pulp ratio[15] and pectoral flipper bone ossification[16]. However, all methods vary in inherent accuracy and must be calibrated against some age estimate of the species in question, most often by using GLG counts, and each have limitations for field use and accuracy across different age classes.

Recent efforts at developing less invasive methods with enhanced accuracy for age determination across multiple species, including marine mammals, has led to a surge in the application of species-specific DNA methylation profiles for the development of epigenetic aging clocks[1, 3, 17–23]. DNA methylation (DNAm) is described as an epigenetic modification whereby a transfer of a methyl (CH_3_) group from S-adenosyl methionine (SAM) to the fifth position of cytosine nucleotides, forming 5-methylcytosine (5mC) nucleotides[24]. The degree of methylation, hypo or hyper can be highly correlated with both chronological age and in the case of abnormal physiological conditions, accelerated aging[25–28].

An epigenetic aging clock was recently published for the bottlenose dolphin (BEAT). The BEAT clock, which relied on methods described for the prior development of a humpback whale specific aging clock, screened 17 CpG sites located within DNA that were harvested from biopsied bottlenose dolphin skin samples to identify two CpGs associated with two genes (*TET2*: CpG site 2, and *GRIA2*: CpG site 5) that have a high correlation (R^2^ = 0.779) with chronological age, error within ± 4.8 years[3, 17]. In addition to the bottlenose dolphin specific BEAT clock, we recently published a multi-tissue Odontocete Epigenetic Aging Clock (OEAC) which accurately (estimates age from blood or skin samples within any odontocete species[1]. The OEAC was developed by applying the same custom DNA Infinium methylation array (HorvathMammalMethylChip40) to skin and blood samples from nine different known age odontocete species, including bottlenose dolphins[1]. Although this odontocete clock works impressively well (median R = 0.95, median absolute error estimation of 2.6 years), evidence suggests that both correlation and accuracy could be improved within each individual species by developing species-specific clocks with an increased sample size for the tissues of interest [29, 30]. For wild bottlenose dolphins, the majority of tissues sampled are collected via remote biopsy and include both skin and blubber samples. Blubber cells are often used to determine hormone concentrations and to evaluate the bioaccumulation of organochloride toxins, while DNA from skin samples is often used to determine relatedness among conspecifics[31–33]. Thus, there is an existing, relatively large historical collection of skin samples and precedent for future collection of skin samples that could also be used for age determination.

In addition to clocks for chronological age determination, recent work in humans has indicated that DNAm clocks can be useful for detecting epigenetic accelerated aging (defined as discrepancy between epigenetic and chronologic age), and that the detection of an accelerated age condition can be predictive of multiple disease states and mortality[34–37]. Epigenetic clocks for bottlenose dolphins may be promising tools for quantifying adverse effects that anthropogenic environmental stressors are creating in wild populations [21].

Therefore, the objectives of our research were 1) to apply the mammalian methylation array technology to develop highly accurate bottlenose dolphin clocks based on blood and/or skin samples; 2) top characterize significant aging associated CpGs identified via epigenome-wide association studies (EWAS); and 3) to characterize sex linked CpGs, and 4) to develop sex estimators for odontocetes based on DNA methylation data.

## Results

We obtained DNA methylation profiles from blood (n = 140) and skin (n = 87) samples from 140 animals (44 males and 96 females) ranging in ages from zero to 57 years of age (mean: 19.8 y, median: 16.8 y). For each animal, we had either blood or skin samples or both. An unsupervised hierarchical analysis clustered the methylation arrays by tissue type (Supplementary Fig. 1).

### Epigenetic aging models

We developed three bottlenose dolphin clocks for i) blood + skin, ii) blood only, and iii) skin only (Fig. 1). To arrive at unbiased estimates of the accuracy, we used Leave One Out Cross Validation (LOOCV). We assessed the correlation between predicted age and chronological age, R, and the median absolute error (MAE). LOOCV reveals that all 3 dolphin clocks are highly accurate: blood and skin clock: R=0.93, MAE = 2.13 years, blood clock: R = 0.97; MAE = 1.46 years, skin clock: R = 0.95; MAE = 2.53 years (Fig 1. b,d,f).

**Fig 1.**
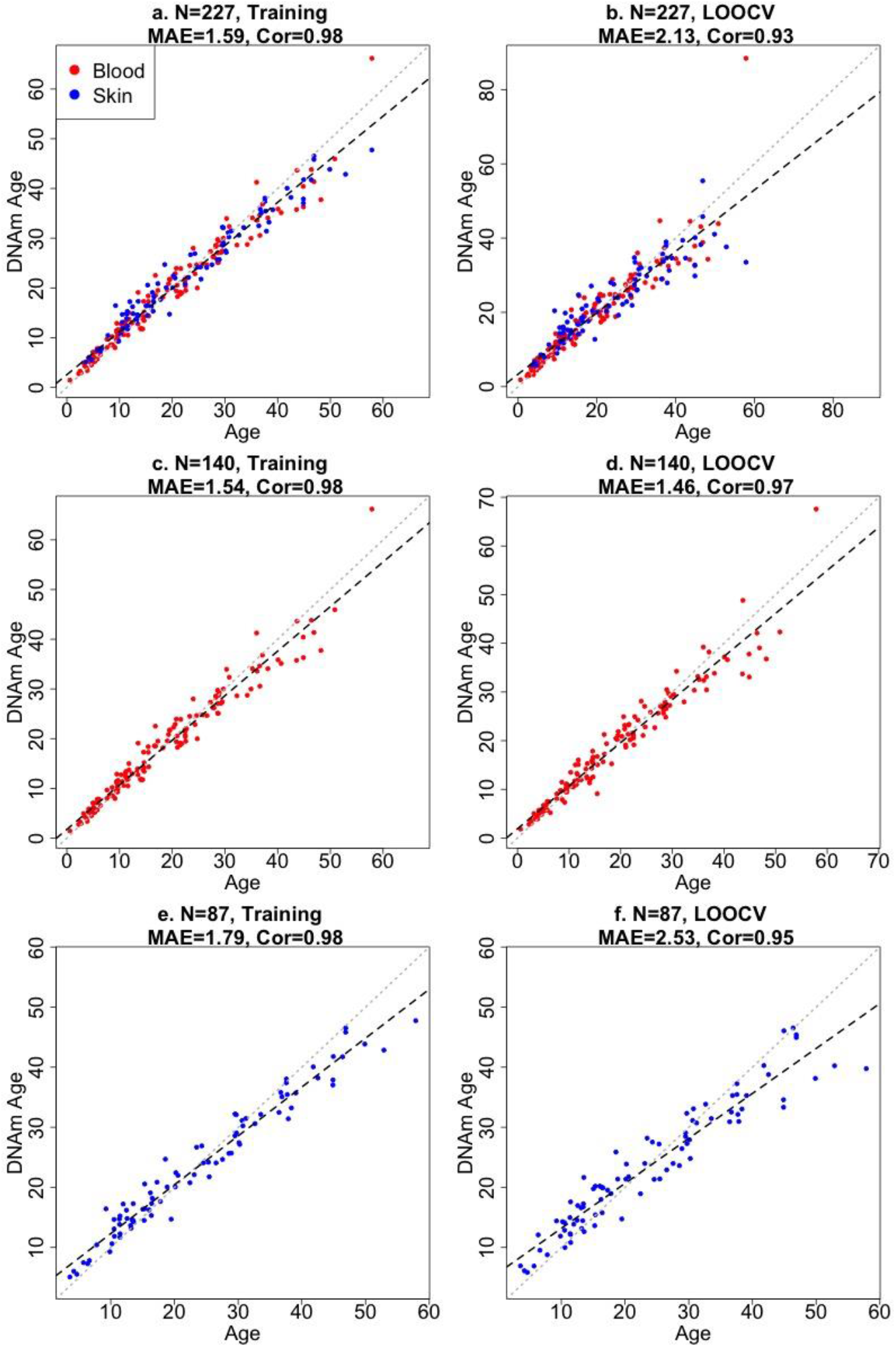
Accuracy of epigenetic clocks for bottlenose dolphin. The panels correspond to three different clocks for bottlenose dolphins: multi tissue clock (a, b); blood clock (c, d); skin clock (e, f) and the age estimates based on training (a, c, e) and LOOCV (b, d, f). Dots are colored by tissue type (red = blood, blue = skin). The training set estimates (left panels) are highly biased. Each panel depicts a linear regression line (black dashed line), a diagonal line (y = x, dotted line), the sample size (N), Pearson correlation (Cor) across all samples, median absolute error (MAE) across species. Chronological age (x-axis) and DNA methylation age estimates (y-axis) are in units of years.

The final versions of the three clocks (based on all training data) involve 48 CpGs (skin and blood clock), 64 CpGs (blood clock), and 39 CpGs (skin clock) as detailed in Supplementary Table 1. The 3 clocks don’t share any CpGs in common (Supplementary Fig. 2). Similarly, the blood clock and the skin clock only share 1 CpG in common (Supplementary Fig. 2).

### Methylation estimator of sex

Methylation based estimators of sex can be useful for identifying human errors in sample labelling or DNA processing. To define a methylation based sex estimator for dolphins and other odontocetes we combined our data with those from 8 additional odontocete species[1]. The sex estimator was developed by fitting a binomial elastic net regression model on n = 447 odontocetes samples (n = 276 female and n = 171 male samples). The model only misclassified two male killer whale samples (99.5% accuracy, Table 2). The 71 CpGs of the sex estimator most likely map to sex chromosomes in odontocetes because they map to human sex chromosomes (based on human genome hg19).

**Table 1.**
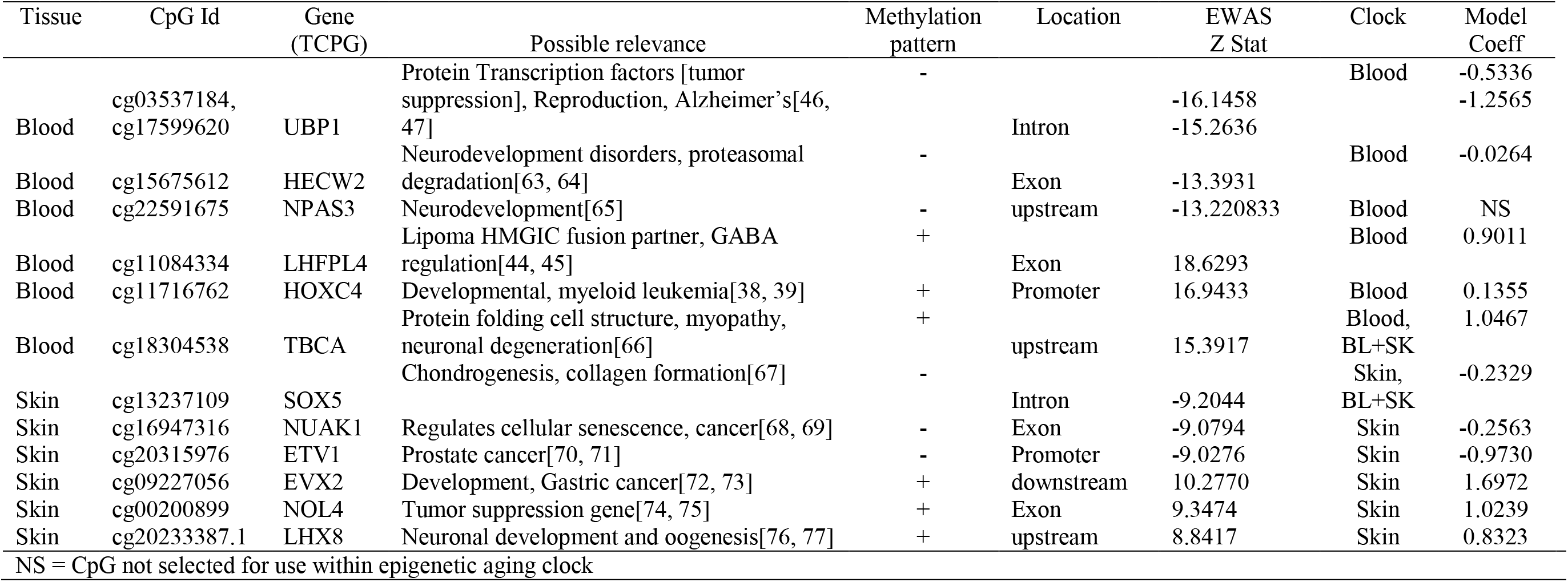
Epigenome-wide associated studies (EWAS) identification of top three hypo [-] and hyper [+] methylated age associated CpGs, its proximate gene, the epigenetic clock which included CpG site its coefficients.

**Table 2.** Elastic Net penalized linear model sex predictions based on odontocetes*, n = 447[1]

|  | Female Samples | Male Samples |
| --- | --- | --- |
| Predicted as Females | 276 | 2 |
| Predicted as Males | 0 | 169 |
| True Positive Rates: | 100 % | 98.83 % |

\*Tissues from odontocete species used in the develop of this sex predictor included: bottlenose dolphin (*Tursiops truncatus*), beluga (*Delphinapterus leucas*), killer whale (*Orcinus orca*). Pacific white-sided dolphin (*Lagenorhynchus obliquidens*), short-finned pilot whales (*Globicephala macrorhynchus*), rough-toothed dolphin (*Steno bredanensis*), Commerson's dolphin (*Cephalorhynchus commersonii*), common dolphin (*Delphinus delphis*), and harbor porpoise (*Phocoena phocoena*).

### Epigenome-Wide Association Studies of age

In total, only 23,005 probes from the mammalian array (HorvathMammalMethylChip40) could be mapped to the bottlenose dolphin genome assembly (turTru1.100). These probes are adjacent to 4558 unique genes: 65% in gene bodies, 10% in promoters, and 25% in distal regions. The number of aligned probes in the bottlenose dolphin is much lower than other cetacean species (e.g., 34,358 probes in the killer whale[1]), which probably reflects the lower quality of the dolphin genome assembly. A total of 6073 and 604 CpGs were significantly correlated with age in blood and skin, respectively (p < 10^−4^, Fig. 2a). This difference in CpG counts between sample types reflects differences in statistical power and sample size (blood, n =140, age range: 1-57 y; skin, n = 87, age range: 4-57 years). The top age-related changes in blood included hypermethylation in *LHFPL4* exon and *HOXC4* promoter. In skin, the CpGs with the strongest positive correlation with age were located downstream of *EVX2* and in an exon of *NOL4* exon (Table 1). CpGs in promoters largely gained methylation with age, which matches DNAm aging pattern in other mammals (Fig. 2b). CpGs located in CpGs islands had higher age correlations than CpGs located outside of islands (Fig. 2c).

**Fig 2.**
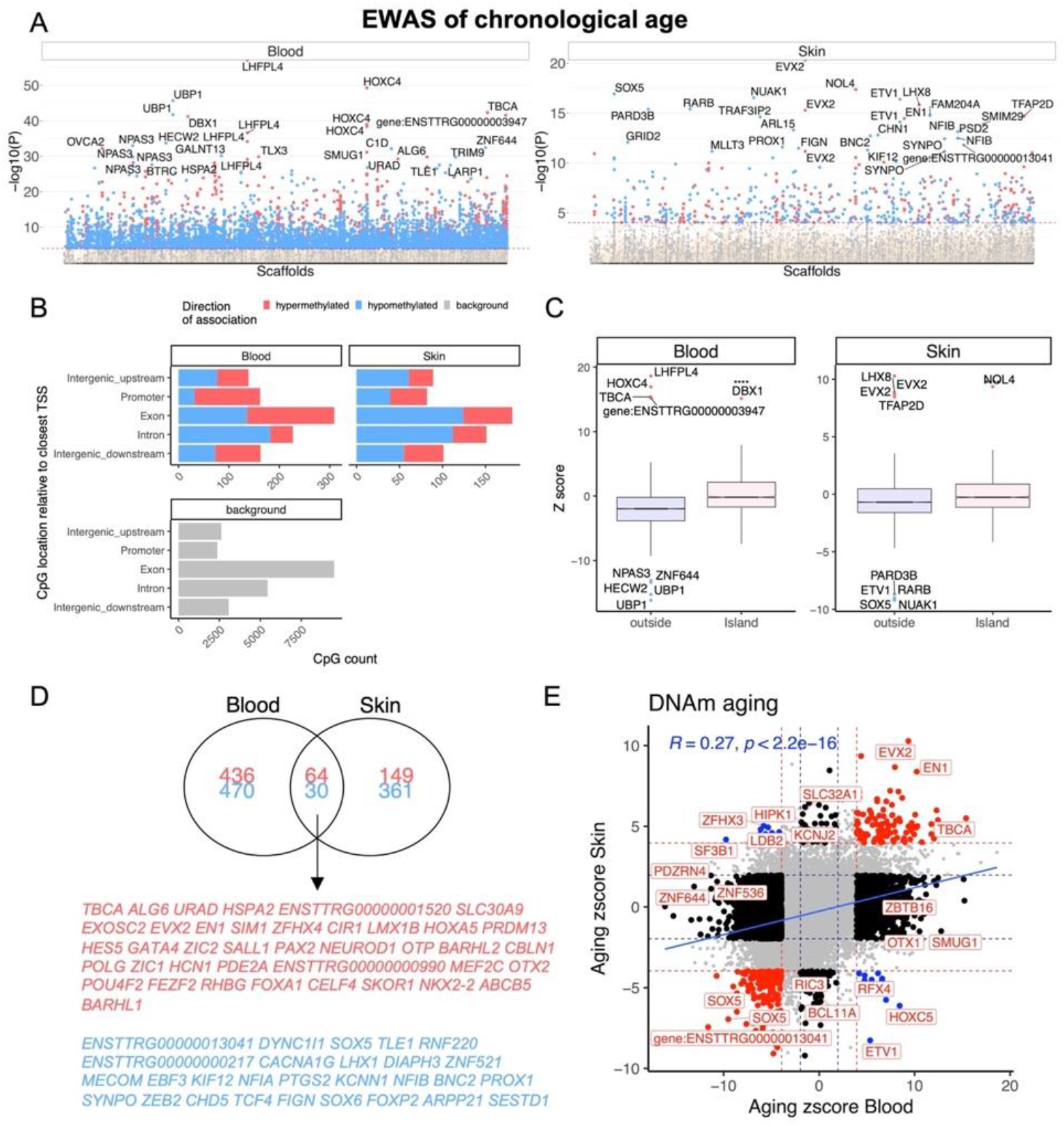
Epigenome-wide association (EWAS) of chronological age in skin and blood of bottlenose dolphin species. (A) Manhattan plots of the EWAS of chronological age. The coordinates are estimated based on the alignment of Mammalian array probes to the bottlenose dolphin (turTru1.100) genome assembly. The direction of associations with P < 0.0001 (red dotted line) is highlighted by red (hypermethylated) and blue (hypomethylated) colors. The 30 most significant CpGs were labeled by their respective neighboring genes. (B) Genomic location analysis of significant (P < 10^−4^) CpGs. The blood analysis focused on the top 500 positively age related CpGs and the top 500 negatively age related CpGs. The skin analysis focused on 213 positively and 391 negatively age related CpGs. (C) Box plot visualizing the median value (line inside the box) and the 25th and 75th percent quartiles (top and bottom of the box), and the whiskers correspond to the 90% percentile. The x-axis corresponds to CpG island status. (D) Venn diagram of the most significant age related CpGs in blood and skin. (E) Sector plot of DNA methylation aging effects in blood and skin. Each axis reports the Z statistic from a correlation test which follows a standard normal distribution under the null hypothesis. Red dotted line: P < 10^−4^; blue dotted line: p > 0.05; Red dots: shared CpGs; black dots: tissue specific changes; blue dots: CpGs where age correlation differs between blood and skin tissue. The grey color in the last panel represents the location of 23,005 mammalian array probes mapped to the bottlenose dolphin (turTru1.100) genome

We found that 94 CpGs (64 hypermethylated, 30 hypomethylated) have a significant correlation with age in both blood and skin (Fig. 2d). Aging effects in blood were moderately correlated with those in skin (R = 0.27, Fig. 2e). Functional enrichment studies reveal that significant age-related CpGs are located near genes that play a role in development and with polycomb repressor complex 2 target genes H3K27ME3, EED, PCR2, and SUZ12 (Supplementary Fig. 3, Table 1). Many CpGs correlate with age in a tissue specific manner. As an example, while HOXC4 promoter, suspected to be involved with mylo-proliferative disorders and skin tumors[38, 39], is hypermethylated with age in blood (z = 4.7), it was hypomethylated in skin (z = −4.5).

### Sex specific DNAm patterns in bottlenose dolphins

Our dataset allowed us to characterize sex differences in baseline methylation levels, and, also, sex effects on aging rates. At a nominal/unadjusted significance threshold of p<10^−4^, 1738 and 742 CpGs were significantly associated with sex in dolphin blood and skin, respectively (Fig. 3a). Due to the incomplete genome assembly we were not able to determine whether these CpGs are located on sex chromosomes in bottlenose dolphins. However, we expect that these CpGs are located on sex chromosomes since more than 70% of these CpGs in blood and 90% in skin were located on the *human* X chromosome. Some of the top sex-related gene regions are as follows: blood, *PTCHD1* exon (hypermethylated in females) and *HUWE1* exon (hypomethylated in females); skin, *HUWE1* exon (hypomethylated in females) and *CNKSR2* exon (hypermethylated in females).

**Fig. 3.**
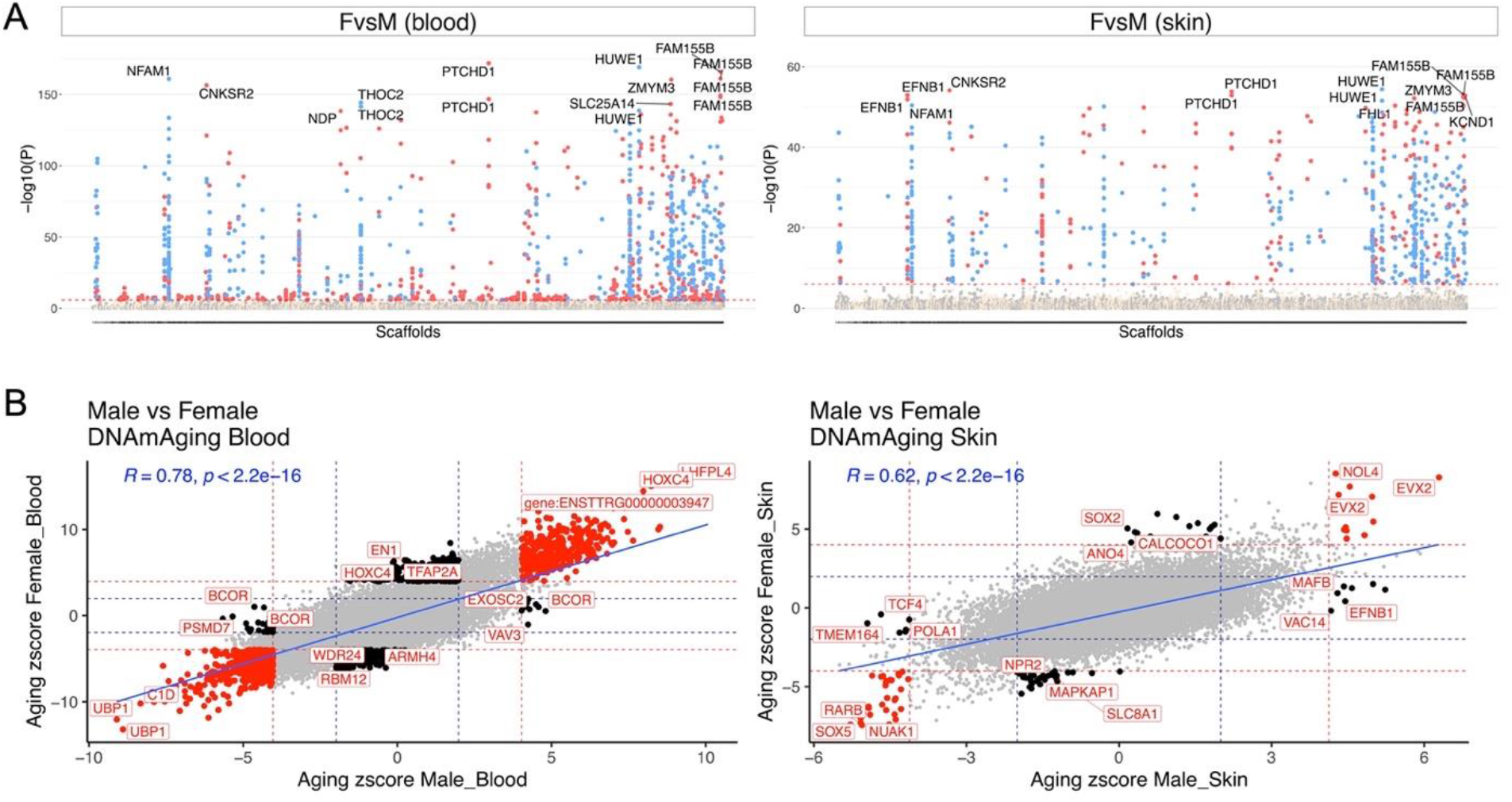
Aging effects in male and female dolphins. A) Manhattan plot of the epigenome-wide association studies (EWAS) of sex. We adjusted the analysis for chronological age. Sample size: males’ blood, 45; females blood, 95; males skin, 29; females skin, 58. The genome coordinates are estimated based on the alignment of Mammalian array probes to the bottle nose dolphin (turTru1.100) genome assembly. The horizontal red dotted line indicate a nominal/uncorrected significance level of P < 10^−4^. Significant CpGs are colored in red if they are hypermethylated in females, blue hypomethylated in females. The 15 most significant CpGs are labeled by their respective neighboring genes. B) Sector plot of DNA methylation aging effects by sex in blood (left) and skin (right). Red dotted line: P < 10^−4^; blue dotted line: P > 0.05; Red dots: shared CpGs; black dots: distinct changes between males and females.

Aging effects were highly correlated between the sexes both in blood (R = 0.78) and skin (R = 0.62, Fig. 3B). We highlight a few CpGs with distinct sex specific aging patterns in Fig. 4.

**Fig. 4.**
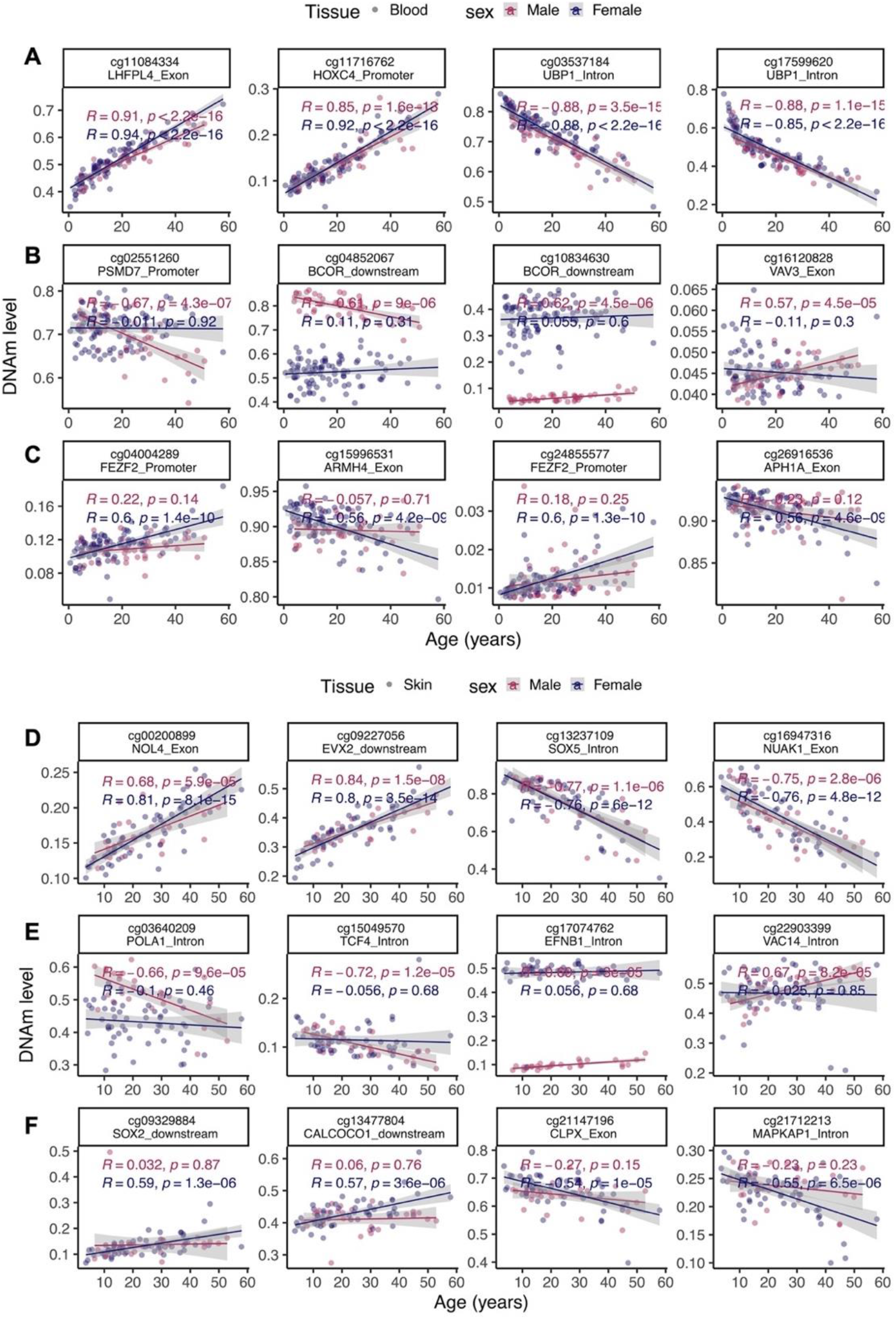
Select age-related CpGs stratified by sex. (A) CpGs that change with age in male and female blood samples from bottlenose dolphins. (B) Male specific changes in blood. (C) Female specific changes in blood. (D) CpGs that change with age in both male and female skin. (E) Male specific changes in skin. (F) Female specific changes in skin. The ordinary least squares regression lines were carried out in males (red) and females (blue) separately.

## Discussion

This study describes the development of a three highly accurate bottlenose dolphin DNAm epigenetic aging clocks using combined blood and skin, blood only or skin only that were developed with samples collected from known (88%) or approximately known age animals. The mammalian methylation array used in this research profiles cytosines that are conserved across mammalian species [1, 22].

Similar to our recently published odontocete epigenetic aging clock (OEAC[1]), the data were mainly comprised of animals whose ages were known precisely (often the birthday was known) or whose age could be estimated with high accuracy. As such, this dolphin clock is expected to become a useful molecular method for accurate age measurement in dolphins.

Compared to recent efforts to develop rapid, non-invasive methods for age determination, including, pectoral flipper bone ossification and dental radiographic determination of the pulp to tooth area ratio which appear to be accurate in juvenile animals[15, 16]. Although bone ossification and pulp to tooth ratios benefit from portability of radiography units that can be used in the field to estimate age within hours of image collection, they still require animal restraint and suffer from reduced accuracy with animals that have reached physical maturity[15, 16]. By contrast, epigenetic clocks apply to the entire life-span of the bottlenose dolphin [1].

Two previous epigenetic clocks apply to Bottlenose dolphins: Bottlenose dolphin Epigenetic Aging estimation Tool (BEAT) and the Odontocete Epigenetic Aging Clock (OEAC)[1, 3]. The OEAC clock is the first multi-tissue clock for multiple odontocetes species. For this clock, nine different odontocete species were used across 446 blood and skin samples, and within this data set, there were 181 bottlenose dolphin samples (140 blood and 41 skin samples). Although we obtained very accurate predictive results for the bottlenose dolphin blood (R = 0.95, MAE 1.5 yr) and skin samples (R = 0.91, MAE 4.8 yr) in our OEAC, predictions based on skin samples were below accuracy levels with a wider variance (MAE) than was found in blood. While still within ranges currently used in forensic sciences for age estimation to within a decade [40–42], this increase in variance in skin samples as compared to blood was hypothesized to be primarily due to a reduced sample size as compared to blood (40 vs 140). Additionally, other variables that may have contributed toward increased predictive variance may have included the types and proportion of cells collected during each skin collections (for discussion see Robeck et al.[1]) and may possibly reflect a higher sensitivity for skin cells to experience varying epigenetic methylation rates across age [43]. Therefore, we hypothesized that by increasing our skin sample size for bottlenose dolphins, we could improve our predictive ability within skin samples over our previous results presented with the OEAC development[1]. Thus, in our present study, we increased our skin sample size by over 100% and obtained an almost a 50% reduction in MAE (2.5 vs 4.8) and brought our median predictive ability from skin samples across all age groups to within 5 years (± 2.5 y). Although these results cannot take away from the accuracy and application of the OEAC across multiple species, they do demonstrate the superior accuracy of species-specific clocks.

Direct comparisons of the accuracy between the BDAC and the OEAC are possible because approximately half of the samples were independently analyzed by each clock. However, direct comparison against the only other bottlenose dolphin clock, the BEAT is not possible due to different samples and measurement platforms. In general, our BDAC skin clock had improved accuracy (R = 0.95, MAE = 2.5 years) at predicting ages of bottlenose dolphins when compared to the BEAT results (BEAT, R^2^ = 0.74, root mean square error = 5.1 years[3]). Important differences between the BDAC and the BEAT clock that have been identified as limitations toward methylation clock development include: (1) the sample number used for the BEAT clock was ∼half (n = 40) the number used for the BDAC; and 2) and the number of CpGs used for final clock development were vastly reduced (2 versus 39). As we have already demonstrated through direct comparisons between the BDAC and OEAC and as was hypothesized by the developers of the BEAT clock, increased sample size has direct effects on clock accuracy[3]. In addition to sample size, by filtering through a dramatically larger catalog of CpGs we were able to identify and incorporate a wider array of significant age related CpG sites into the predictive model resulting an improved representation of organism age [22, 29, 30].

The BEAT clock [3] identified two CpG sites on two genes, *TET2* (CpG site 2) and *GRIA2* (CpG site 5) that accounted for 78% of the age-related variation in % DNA methylation. These same genes were also found to correlate with age in humpback whales[17]. Although we had several probes that mapped to the genomic regions of *GRIA2*, this gene was not considered to be within the top 10 genes that accounted for age-related methylation changes in neither our present study nor within the OEAC[1]. Our measurement platform (the mammalian array) did not cover the *TET2* gene, and we only had probes that mapped to the genomic regions of *GRIA2* gene. Interestingly, most of these *GRIA2* probes (both exon and intron) were hypomethylated with age in both the blood and skin of bottlenose dolphins (Supplementary Fig. 4). This suggests that although promoter of *GRIA2* gains methylation with age[3], selected CpGs in the gene body lose methylation with age.

Besides epigenetic clock development, the mammalian array is a uniquely reproducible tool for a direct genome-wide comparison of DNAm changes across cetaceans. Our EWAS identified DNAm aging CpGs proximate to genes associated with development (Supplementary Fig. 3). The top proximate hypermethylated CpG in blood was associated with *LHFPL4*, a gene known to be involved with GABA regulation in the central nervous system and peripheral lipoma formation[44, 45]. This gene has recently been identified as the most predictive age-related gene across mammalian species and despite its conserved status, its contribution towards age-related functional decline has not be clarified[37]. The top two hypomethylated CpGs (both within introns) in the blood were proximal to *UBP1*, a gene that is closely related to *TFCP2* and is considered part of the Grainyhead family of transcription factors[46]. Although *UBPI* has been identified as involved in angiogenesis during growth and development and Alzheimer’s disease, its redundancy and homology with *TFCP2* make its yet unidentified role in cancer regulation a likely possibility[46, 47]. For skin, the top proximate genes were associated with development and energy metabolism (Supplementary Fig. 3). In addition to these CpG related genes, multiple CpGs, in both blood and skin, that were associated with genes within the PRC2 complex were hypermethylated (Supplementary Fig. 3). The PRC2 is required for embryonic stem cell differentiation but also plays a role in in aging [48, 49]. Overall, we find that age-related methylation changes in bottlenose dolphins mirror those in other odontocete species and mammals in general [1, 37].

Our sex related CpGs were largely the same as those found in multiple odontocete species [1]. Therefore, we were able to use these CpGs to develop a highly accurate (99.5%) multivariate predictor of sex across all odontocete species. Although not of primary importance for this research, the ability to determine sex in species that do not have easily identifiable sexual dimorphic patterns is an attractive property of cytosine methylation data.

Similar to previous work with other cetaceans, we found no evidence that accounting for sex was necessary for the development of the three BDAC clocks[1, 23]. Aging effects in males were highly correlated with those in females both in blood (R = 0.78) and skin (R = 0.62). While the majority of sex specific CpGs were found on the X chromosome (78.8%, Supplementary Data File 1), the proximate genes associated with the top non X chromosome CpGs, which were different between the two sexes included in blood, related to cancer (*PSMD7*[50], *BCOR*[50]) and the central nervous system (*FEZF2* [51]). Noteworthy sex related autosomal genes implicated by the skin methylation data play a role in DNA repair (*POLA1*[52]), neurodevelopment (*TCF4*[53]), oral cancer (SOX2[54], higher methylation rates in females) and nutritional or chemical mediated ER autophagy (CALCOCO1[55]). These results suggest a potential difference in aging phenotypes in dolphin sexes, however, this hypothesis should be examined in future studies.

In conclusion, we characterized cytosines that correlate strongly with age and sex in bottlenose dolphins. We present three highly accurate DNA methylation-based estimators of chronological age for bottlenose dolphins. The high accuracy of these clocks could not have been achieved without knowing the precise ages of a large number of animals which in turn relied on long term efforts of monitoring and caring for these animals.

## Materials and Methods

### Ethics approval

The study was authorized by the SeaWorld Parks and Entertainment and Miami Seaquarium animal care and use committee.

### Study Animals

For model development, our study population included skin (n = 50) and blood samples (n = 140) from 140 bottlenose dolphins located at three SeaWorld Parks (Orlando, San Antonio, and San Diego) and Discovery Cove (Orlando, FL). The known age animals consisted of 123 zoo born animals (38 male, 85 females) with a median age of 14.5 y (range: 0.57 to 40.7 y), and 17 wild born animal (6 male, 11 females) with a median age of 36.4 years range: 3.9 to 57.0 y). Known (87.9%) or estimated (based on length at capture or rescue for stranded animals) birth dates were used for correlating against methylation predicted age

### Sample collection

Blood samples were collected either voluntarily from the peripheral periarterial venous rete on the ventral tail fluke using an 18 to 22 gauge winged blood collection set or attached to a vacutainer collection system during routine physical examination. Blood was collected by either the veterinary technician or staff veterinarian into BD Vacutainers (Becton Dickinson, Franklin Lakes, NJ) containing EDTA. Samples were inverted in the Vacutainer a minimum of 10 times and then frozen at −80°C until further testing.

Skin scrapings were collected either under stimulus control or manual restraint using a sterile disposable dermal curette or 8.0 mm biopsy punch (Miltex, Integra Life Sciences Corp.,York, PA) from a location just posterolateral of the dorsal fin overlying the epaxial muscle. Prior to collection, a cold pack was placed on the site for several minutes prior to sampling to numb the sample site. Skin samples were placed into sterile cryovials (Nunc® Cryotubes, Millipore Sigma Corp., St. Louis, MO) and stored at −80°C until shipment on dry ice. Skin samples from non-living animals were obtained from frozen (−80°C) specimens that had been previously collected and stored during standard necropsy procedures.

### DNA extraction

Genomic DNA was extracted from clotted whole blood samples using QIAamp DNA Mini blood kit and following the manufacturer’s instructions (Qiagen, Valencia, CA). Tissue samples were pulverized and broken down manually using a drill and DNA was extracted using DNeasy Tissue kit (Qiagen) and following the manufacturer’s instructions with the exception of extending the proteinase k digestion. DNA was then extracted using the automated nucleic acid extraction platform, Anaprep (Biochain, Newark, CA) that utilizes a magnetic bead extraction process and Tissue DNA Extraction kit (Anaprep).

### DNA methylation data

The mammalian DNA methylation arrays were profiled using a custom Infinium methylation array (HorvathMammalMethylChip40) based on 37,491 CpG sites as previously described[56]. Out of these sites, 1951 were selected based on their utility for *human* biomarker studies; these CpGs, which were previously implemented in human Illumina Infinium arrays (EPIC, 450K), were selected due to their relevance for estimating age, blood cell counts or the proportion of neurons in brain tissue. The remaining 35,540 probes were chosen to assess cytosine DNA methylation levels in mammalian species[57]. The subset of species for each probe is provided in the chip manifest file at the NCBI Gene Expression Omnibus (GEO) platform (GPL28271). Raw data was normalization using the SeSaMe method which assigned a beta value (0 to 1) for each methylation estimate corresponding to each probe used for every sample[58]. Beta values were indicative of methylation rates, with 0 indicating no gene copies were methylated. Unsupervised hierarchical clustering analysis based on the interarray correlation was used to identify technical outliers which were then removed from further analysis.

### Penalized regression

Details on the clocks (CpGs, genome coordinates) and R software code are provided in the Supporting (SI) Appendix file. Penalized regression models were implemented with the R software package “glmnet“[59]. The optimal penalty parameters lambda in all cases were determined automatically by using a 10 fold internal cross-validation (cv.glmnet) on the training set. Alpha=1/2 corresponds to the elastic net penalty that penalizes the coefficients based on their magnitude. We performed a Leave One Sample Out Cross Validation (LOOCV) scheme for arriving at unbiased estimates of the prediction accuracy of the different DNAm aging clocks. The LOOCV, which is based on previously reported methods[60, 61], does the following for each of the *N* samples: omit the one sample from the training set; fit the clock on the training set with (*N* - 1) samples; predict the DNAm age of the omitted sample with the fitted clock. Therefore, the LOOCV allows one to estimate the accuracy of predicting the age for any unknown bottlenose dolphin sample by the clock. The cross-validation study reports unbiased estimates of the age correlation r (defined as Pearson correlation between the DNAm age estimate and chronological age) as well as the median absolute error (MAE). The accuracy of the resulting bottlenose dolphin clocks was assessed via LOOCV estimates of: 1) the correlation r between the predicted epigenetic age and the actual (chronological) age of the animal; and 2) the median absolute error (MAE) between DNAm age and chronological age.

### Multivariate sex predictor

To predict sex (binary variable with values 1=female, 0=male), we used a binomial generalized linear model, regularized by elastic net[62]. The glmnet alpha parameter was set to 0.5, i.e. we used elastic net regression. The lambda parameter of the glmnet function was chosen using 10-fold cross validation. As we observed a reasonably balanced female-male sample sizes in the data set, we set a neutral prediction rule for the binomial generalized linear model. If the predicted probability for a sample being female was greater than 0.5, the sample was predicted to be female. We used all odontocetes samples previously used for DNAm clock develeopment[1] and the additional samples from bottlenose dolphins, reported herein, in the training data so that the resulting sex estimator will be broadly applicable to odontocetes species.

## Data availability

The data will be made publicly available as part of the data release from the Mammalian Methylation Consortium. Genome annotations of these CpGs can be found on Github https://github.com/shorvath/MammalianMethylationConsortium.

## Code availability

The R software code used for clock development are provided in the Supplement.

## Acknowledgements

We thank individuals from all of our zoological parks that contributed to toward sample collection and curating for this research. Specifically, from SeaWorld California (SWC) we thank Dr. Elsburgh Clarke, Dr. Kelsey Herrick, Rachelle Pastorkovich, Sarah McMillen, Melinda Tucker, Kim Regan, and Wendy Ramirez. From SeaWorld of Texas, we thank Dr. Jennifer Russel and Tara Klimek. From SeaWorld of Florida, we thank Dr. Dana Lindemann, Dr. Claire Erlacher-Reid, Jacob Vandenberg, Cynthia O’neil, Megan Fox, Siobhan Diaz, Cynthia Reyes, Stephanie Smith and Heidy Clifford. Finally, we thank Species Preservation Laboratory research technicians Amanda McDonnel and Jacqueline Posy. This is a SeaWorld Parks and Entertainment technical manuscript number 2021-7.

