## Supplementary Figures and Tables for "Multi-tissue methylation clocks for age estimation in the common bottlenose dolphin"

*Steve Horvath, PhD, ScD*

*Todd Robeck, PhD, DVM*

**

**This PDF file includes**

**Supplementary Methods**

**Supplementary Table 1**

**Supplementary Figures S1 to S6**

**Supplementary Methods**

**Statistical details for epigenetic clock development.**

All of the bottlenose dolphin (*Tursiops truncatus*), blood+skin, blood and skin clocks use the same age transformation. Denote sample $i$ from species $k$ has Age $X_{ik}$ and Age at sexual maturity for species $k$ is $a_{k}$, Gestation time (in years) for species $k$ is $g_{k}$. Then the log-linear transformation is as follows (Equation 1)

$$Equation 1. Y_{ik}=\left\{ \begin{matrix} \log(\frac{X_{ik}+g_{k}}{1.5a_{k}+g_{k}}), & X_{ik}\leq1.5a_{k} \\ \frac{X_{ik}-1.5a_{k}}{1.5a_{k}+g_{k}}, & X_{ik}>1.5a_{k}. \end{matrix} \right.$$

Adding gestation time in the transformation means age since inception and avoids negative values in the $\log$. This transformation ensures the transformed age $Y_{ik}$ is continuous and has continuous first derivative at $X_{ik}=1.5a_{k}$.

The Bottlenose dolphin clocks use an age transformation that is dependent on a mean age between male and female at sexual maturity of 8.93 years and a gestation time of 1.03 years. 1

| **Supplementary Table 1**. The CpG (Cg) sites selected for use within each bottlenose dolphin multi-tissue and tissue specific epigenetic clocks. The CpG sites are listed with the main categorical location (intron, exon or promotor etc.) and proximate gene within SI Data File 1. | | | | | |
| --- | --- | --- | --- | --- | --- |
| **BD Blood & Skin Clock** | | **BD Blood Clock** | | **BD Skin Clock** | |
| **CpG site** | **Coefficient** | **CpG site** | **Coefficient** | **CpG site** | **Coefficient** |
| (Intercept) | 5.290 | (Intercept) | 0.613 | (Intercept) | 2.932 |
| cg00200899 | 0.354 | cg00301020 | -0.047 | cg00200899 | 1.024 |
| cg00473176 | -0.542 | cg00801362 | -0.220 | cg01893274 | -0.023 |
| cg00493831 | 1.791 | cg01965931 | 0.001 | cg03225697 | -0.345 |
| cg01893274 | -0.444 | cg03524277 | -0.701 | cg05111574 | -0.321 |
| cg02317025 | 0.742 | cg03537184 | -0.534 | cg07010480 | -1.208 |
| cg02334175 | -1.284 | cg04184175 | -0.332 | cg08125125 | -0.209 |
| cg04041705 | -0.268 | cg04633983 | 0.169 | cg08796632 | -0.001 |
| cg04184175 | -1.092 | cg05584808 | 0.799 | cg08815701 | -0.187 |
| cg05463929 | 0.855 | cg06569016 | 0.507 | cg09227056 | 1.697 |
| cg05584808 | 0.458 | cg06742640 | -0.003 | cg09817427 | 1.409 |
| cg05991926 | -0.340 | cg06750335 | -0.054 | cg10501210 | -0.114 |
| cg07366921 | -1.712 | cg06836529 | -0.113 | cg10620475 | -0.050 |
| cg07544506 | -1.520 | cg07213500 | 0.002 | cg10763467 | 0.392 |
| cg08563010 | -0.269 | cg09296085 | -0.024 | cg10876801 | -0.236 |
| cg09363187 | 0.145 | cg10501210 | -0.351 | cg12905991 | 0.233 |
| cg09497746 | 3.569 | cg10504143 | 1.514 | cg13237109 | -0.233 |
| cg10504143 | 0.429 | cg10749945 | 0.052 | cg13400013 | 0.729 |
| cg10975089 | -0.353 | cg10918171 | 0.007 | cg13837730 | -0.050 |
| cg11622231 | 0.093 | cg11084334 | 0.901 | cg15661483 | -0.266 |
| cg11723266 | -0.586 | cg11104864 | -0.100 | cg15809488 | -0.082 |
| cg11857072 | -0.120 | cg11622231 | 0.003 | cg15974234 | -0.158 |
| cg12540262 | -0.144 | cg11716762 | 0.136 | cg15976167 | -0.155 |
| cg12763898 | 0.623 | cg11723266 | -0.272 | cg16947316 | -0.256 |
| cg12807727 | -0.238 | cg11728741 | 2.030 | cg17829420 | -0.532 |
| cg12841266 | 1.893 | cg12540262 | -0.034 | cg17856858 | -0.480 |
| cg13237109 | -0.569 | cg12841266 | 0.702 | cg17983750 | -0.030 |
| cg13400013 | 1.734 | cg12879445 | 0.110 | cg18433728 | -0.145 |
| cg13433278 | 0.656 | cg12946225 | 0.142 | cg18889628 | -0.029 |
| cg13502730 | 0.223 | cg13367380 | 0.135 | cg19565468 | 0.482 |
| cg15974234 | -0.325 | cg13502730 | 0.127 | cg20233387.1 | 0.832 |
| cg15976167 | -0.120 | cg14115575 | 0.392 | cg20315976 | -0.973 |
| cg17437489 | -0.394 | cg14182735 | -0.281 | cg21477760 | -0.144 |
| cg17594003.1 | -0.698 | cg15675176 | 0.151 | cg21628661 | -0.199 |
| cg18039160 | -0.105 | cg15675612 | -0.026 | cg23360751 | -0.171 |
| cg18092286 | -0.225 | cg17437489 | -0.188 | cg24104506 | 0.328 |
| cg18304538 | 6.536 | cg17594003.1 | -0.109 | cg24615697 | -0.136 |
| cg19489331 | -0.665 | cg17599620 | -1.256 | cg26535952 | -0.681 |
| cg19834279 | -0.315 | cg17919530 | -0.425 | cg27245416 | -0.021 |
| cg19981759 | 0.226 | cg17997082 | 0.170 |  |  |
| cg23449696 | 0.558 | cg18278920 | 0.786 |  |  |
| cg23576695 | -0.002 | cg18304538 | 1.047 |  |  |
| cg25430089 | -0.638 | cg18473521.1 | 0.041 |  |  |
| cg26062663 | 1.420 | cg18473521.2 | 0.135 |  |  |
| cg26679061 | 1.250 | cg18659931 | -0.233 |  |  |
| cg26931581 | 3.720 | cg19204561 | -0.717 |  |  |
| cg27201382 | 1.632 | cg19868479 | -0.091 |  |  |
| cg27557124 | -0.133 | cg19965683 | -0.070 |  |  |
|  |  | cg20142163 | 0.048 |  |  |
|  |  | cg20473688 | -0.011 |  |  |
|  |  | cg20699548 | -0.108 |  |  |
|  |  | cg21097283 | 0.130 |  |  |
|  |  | cg23004138 | 0.082 |  |  |
|  |  | cg23087015 | 0.146 |  |  |
|  |  | cg23931487 | 0.003 |  |  |
|  |  | cg24015091 | 0.072 |  |  |
|  |  | cg24805210 | -0.279 |  |  |
|  |  | cg24866418 | 0.007 |  |  |
|  |  | cg25184464 | -0.014 |  |  |
|  |  | cg25430089 | -0.093 |  |  |
|  |  | cg26554534 | -0.340 |  |  |
|  |  | cg26957053 | 0.072 |  |  |
|  |  | cg26979561 | -0.026 |  |  |
|  |  | cg27088374 | -0.089 |  |  |

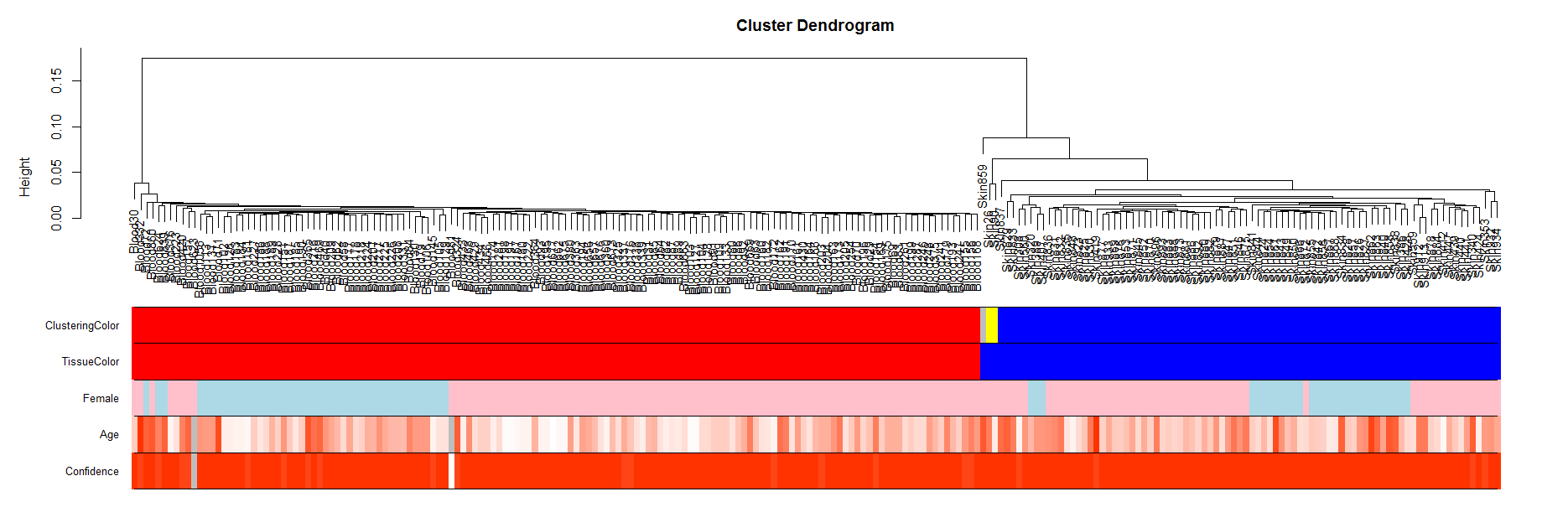

**Supplementary Fig. 1 Unsupervised hierarchical clustering of tissue samples from dolphins**. Average linkage hierarchical clustering based on the interarray correlation coefficient (Pearson correlation). The cluster branch (first color band) corresponds to tissue type (second color band). Third color band encodes sex (pink=female). Fourth color band encodes age (red is a high age, white low age). Final color band encodes the confidence in the age (90 percent or higher for most samples).

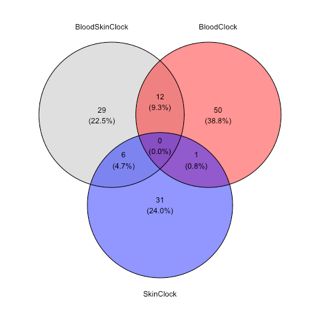

**Supplementary Fig. 2. Shared numbers of CpGs of the 3 different dolpin clocks.** The 3 clocks don't share any CpGs in common. The blood clock and the skin clock only share 1 CpG in common.

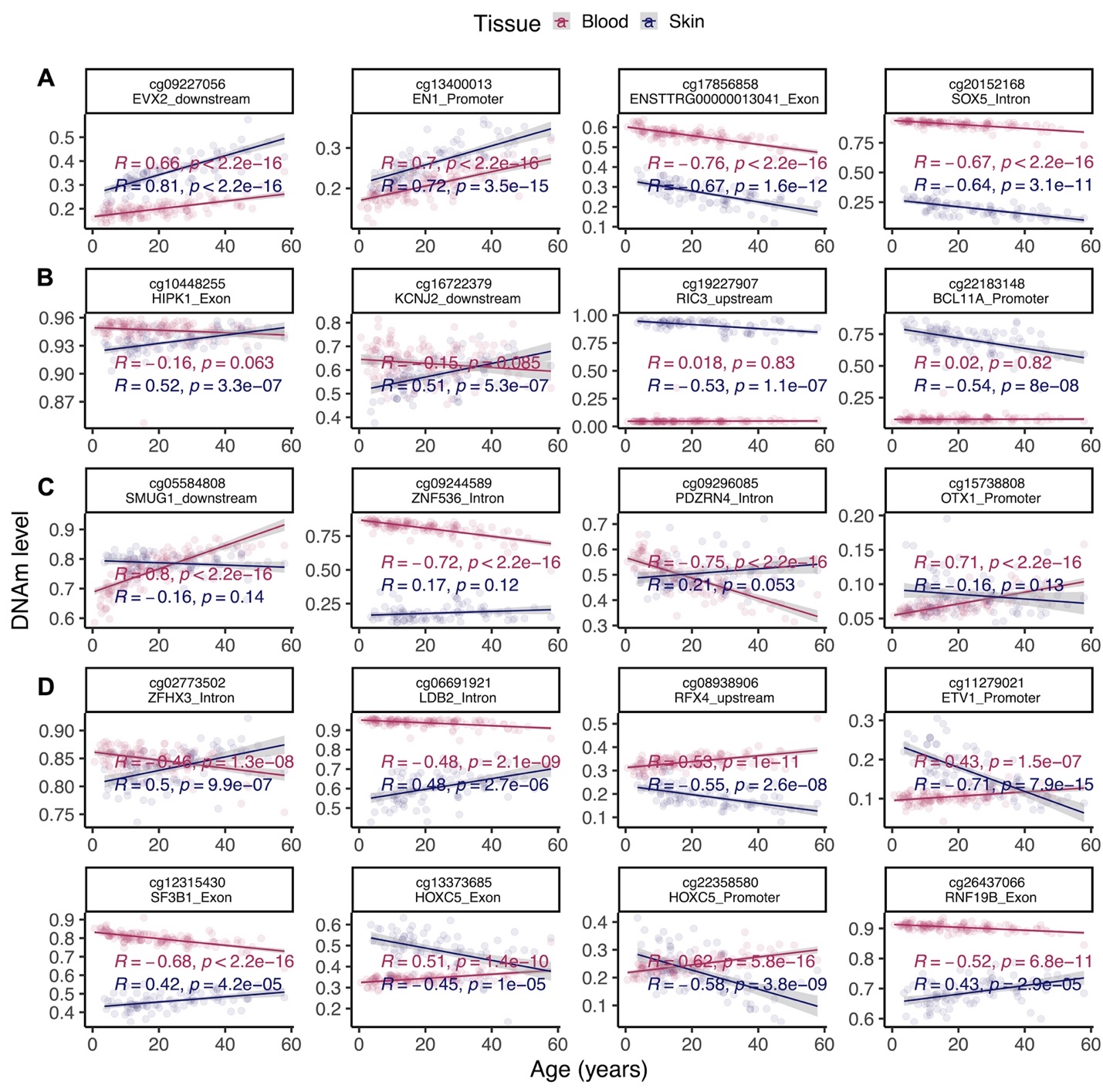

**Supplementary Fig. 3. Scatter plots of selected CpGs that change with age in bottlenose dolphins.** (A) CpGs that change with age in both blood and skin. (B) Skin specific changes. (C) Blood specific changes. (D) Example CpGs with divergent DNAm aging between blood and skin.

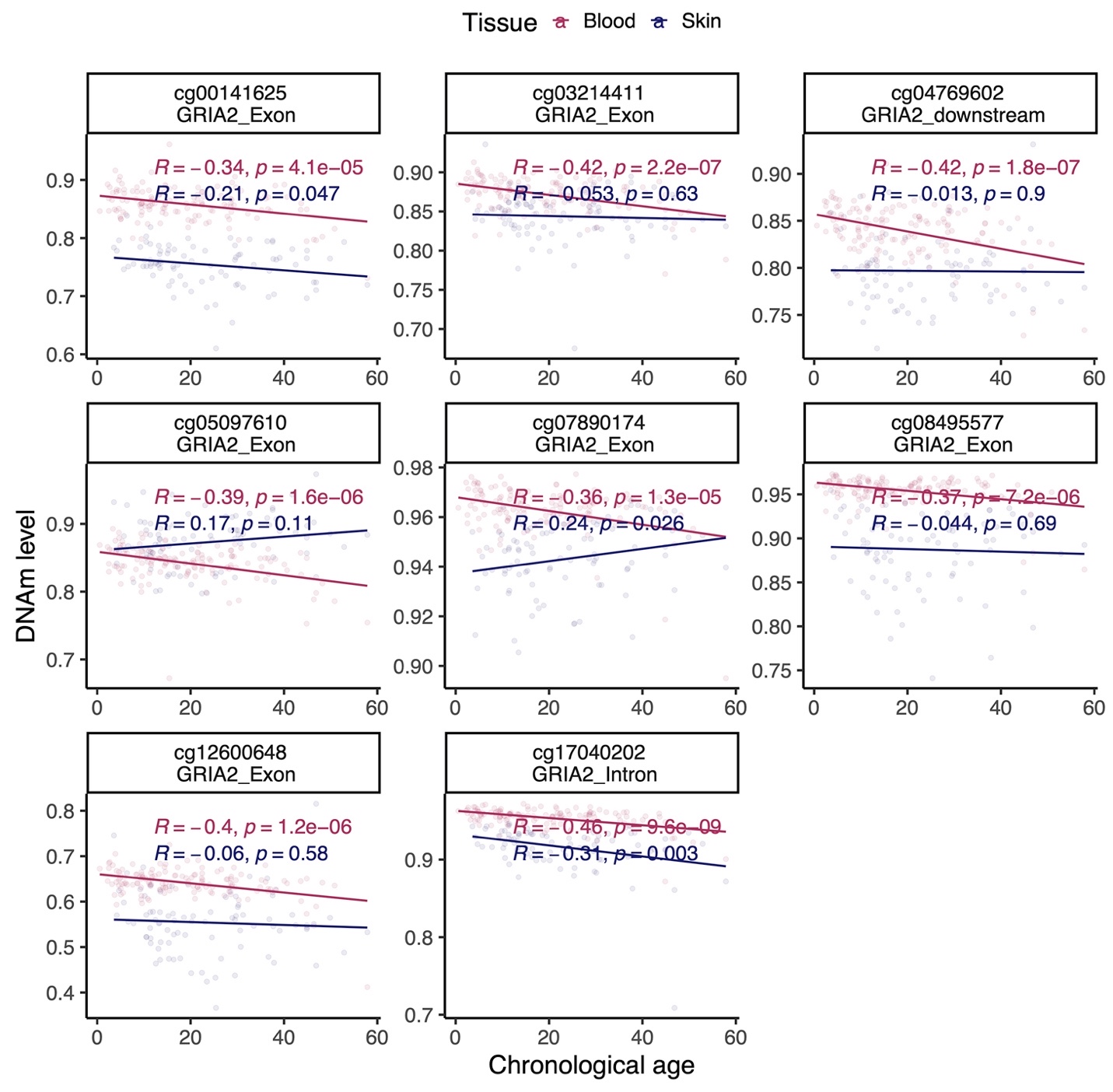

**Supplementary Fig. 4.** **Scatter plots of CpGs adjacent to GRIA2 gene that change with age in bottlenose dolphins.**

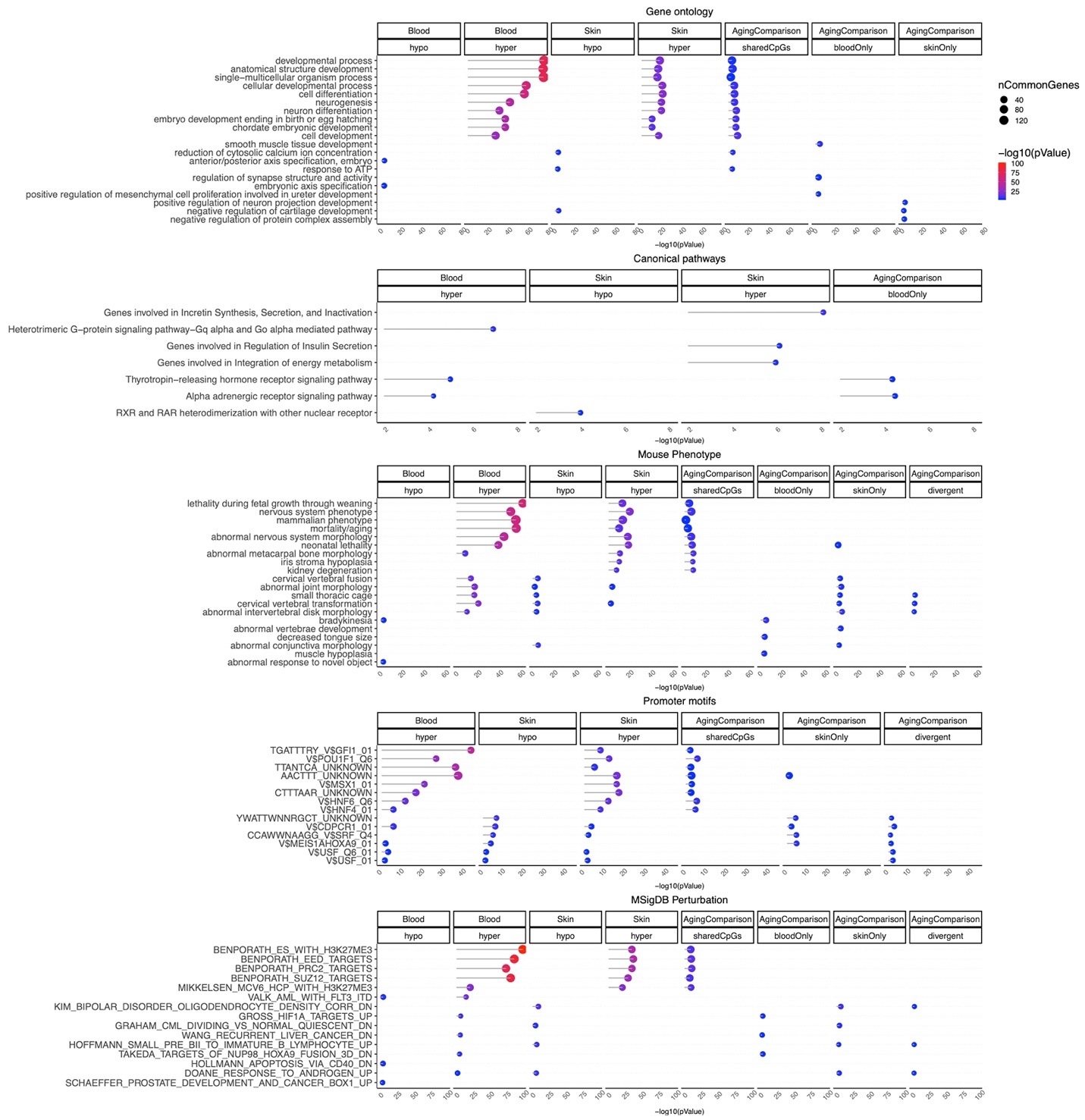

**Supplementary Fig. 5.** **Gene set enrichment analysis of DNAm aging blood and skin of bottlenose dolphins.** The gene level enrichment was done using GREAT analysis and human Hg19 background limited to the probes that mapped to turTru1.100 genome. Datasets: gene ontology, canonical pathways, mouse phenotypes, promoter motifs, and MSigDB Perturbation. The results were filtered for significance at p < 10^-3^. Aging comparison columns are the CpGs that defined based on the sector plot in Figure 4D.

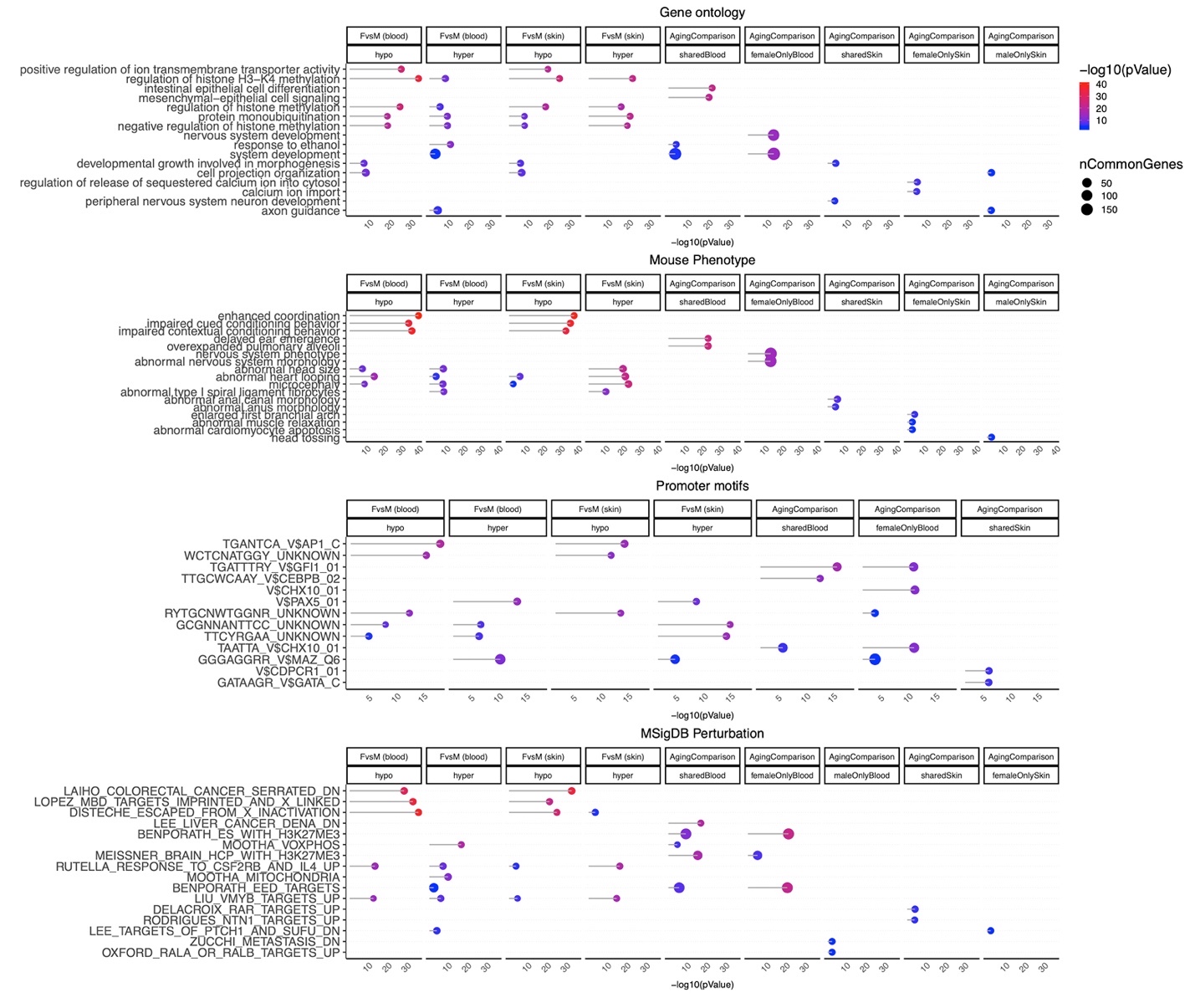

**Supplementary Fig. 6.** **Gene set enrichment analysis of tissue specific sex differences in bottlenose dolphins.** The gene level enrichment was done using GREAT analysis and human Hg19 background limited to the probes that mapped to turTru1.100 genome. Datasets: gene ontology, mouse phenotypes, promoter motifs, and MSigDB Perturbation. The results were filtered for significance at p < 10^-3^. “FvsM” columns are the CpGs with basal (mean) methylation difference between sexes regardless of chronological age of the animals. Aging comparison columns are the CpGs that defined based on the pattern of DNAm aging in sector plots in Figure 5B.
